# Alignment by numbers: sequence assembly using compressed numerical representations

**DOI:** 10.1101/011940

**Authors:** Avraam Tapinos, Bede Constantinides, Douglas B Kell, David L Robertson

## Abstract

**Motivation:** DNA sequencing instruments are enabling genomic analyses of unprecedented scope and scale, widening the gap between our abilities to generate and interpret sequence data. Established methods for computational sequence analysis generally use nucleotide-level resolution of sequences, and while such approaches can be very accurate, increasingly ambitious and data-intensive analyses are rendering them impractical for applications such as genome and metagenome assembly. Comparable analytical challenges are encountered in other data-intensive fields involving sequential data, such as signal processing, in which dimensionality reduction methods are routinely used to reduce the computational burden of analyses. We therefore seek to address the question of whether it is possible to improve the efficiency of sequence alignment by applying dimensionality reduction methods to numerically represented nucleotide sequences.

**Results:** To explore the applicability of signal transformation and dimensionality reduction methods to sequence assembly, we implemented a short read aligner and evaluated its performance against simulated high diversity viral sequences alongside four existing aligners. Using our sequence transformation and feature selection approach, alignment time was reduced by up to 14-fold compared to uncompressed sequences and without reducing alignment accuracy. Despite using highly compressed sequence transformations, our implementation yielded alignments of similar overall accuracy to existing aligners, outperforming all other tools tested at high levels of sequence variation. Our approach was also applied to the *de novo* assembly of a simulated diverse viral population. Our results demonstrate that full sequence resolution is not a prerequisite of accurate sequence alignment and that analytical performance can be retained and even enhanced through appropriate dimensionality reduction of sequences.

## 1 INTRODUCTION

Contemporary sequencing technologies have massively parallelised the determination of nucleotide order within genetic material, making it possible to rapidly sequence the genomes of individuals, populations and interspecies samples (Bentley et al., 2008; Eid et al., 2009; Margulies et al., 2005; Rothberg et al., 2011; Salipante et al., 2014). However, the sequences generated by these instruments are usually considerably shorter in length than the genomic regions they are used to study. Genomic analyses accordingly begin with the process of sequence assembly, wherein sequence fragments (reads) are reconstructed into the larger sequences from which they originated. Computational methods play a vital role in the assembly of short reads, and a variety of assemblers and related tools have been developed in tandem with emerging sequencing platforms (Bradnam et al., 2013; Schatz et al., 2010; Shendure and Aiden, 2012).

Where the objective of a nucleotide sequencing experiment is to derive a single consensus sequence representing the genome of an individual, various computational methods are applicable. Seed-and-extend alignment methods using suffix array derivatives such as the Burrows-Wheeler Transform have emerged as the preferred approach for assembling short reads informed by a supplied reference sequence (Li and Durbin, 2009; Shrestha et al., 2014), while graph-based methods employing Overlap Layout Consensus (OLC) (Kececioglu and Myers, 1995; Myers, 1995) and de Bruijn graphs of *k*-mers (Earl et al., 2011; Iqbal et al., 2012; Pevzner et al., 2001) have become established for reference-free *de novo* sequence assembly. However, the effectiveness of these approaches varies considerably (Bradnam et al., 2013) for characterising genetic variation within populations (*‘deep’* sequencing), or inter-species biodiversity within a metagenomic sample.

Within populations comprising divergent variants, such as those established by a virus within their host, bias associated with the use of a reference sequence can lead to valuable read information being discarded during assembly (Archer et al., 2010). While this can be overcome by constructing a data-specific reference sequence after initial reference alignment, this still necessitates use of a reference sequence for the initial alignment, a limiting factor for sequencing unknown species, often abundant in metagenomic samples. On the other hand, while *de novo* approaches require little *a priori* knowledge of target sequence composition, they are very computationally intensive and their performance scales poorly with datasets of increasing size (Myers, 1995). Indeed, the underlying problem of *de novo* assembly is NP-hard, with the presence of genetic diversity serving to increase further the complexity of computational solutions. As such, aggressive heuristics are typically employed in order to reduce the running time of *de novo* assemblers, which in turn can compromise assembly quality.

The difficulty of diverse population assembly is exacerbated by the scale of sequencing datasets. Contemporary sequencing platforms may generate billions of reads per run, posing considerable challenges for timely computational sequence analysis. Of these, a key challenge to be overcome when analysing any sufficiently large dataset arises when its size exceeds the capacity of a computer’s working memory. If, for example, stored short read data exceeds the capacity of a computer’s random access memory (RAM), these data must be exhaustively ‘swapped’ between RAM and disk storage that is orders of magnitude slower to access, forming a major analysis bottleneck (Yang and Wu, 2006). *de novo* assembly in particular requires manyfold more memory, *O(n*^*2*^),than is needed to store the read information itself. Additionally, all-to-all pairwise comparison of reads scales poorly (quadratic *O(n*^*2*^)time complexity) with increasing data size. While indexing structures such as suffix arrays are often used to reduce the burden of pairwise sequence comparison, *O(n log(n))*, their performance generally deteriorates with increasing sequence length, in accordance with the phenomenon known as the ‘curse of dimensionality’ (Verleysen and François, 2005).

Comparable analytical challenges involving high dimensional sequential data are encountered in other data-intensive fields such as signal and image processing, and time series analysis, where a number of effective dimensionality reduction methods have been proposed, including the discrete Fourier transform (DFT) (Agrawal et al., 1993), the discrete wavelet transform (DWT) (Chan and Fu, 1999; Woodward et al., 2004), and piecewise aggregate approximation (PAA) (Geurts, 2001; Keogh et al., 2001). The DFT or DWT may be used to transform data into their frequency or wavelet domains respectively, enabling the identification of major data characteristics and the suppression of minor features and/or noise (Shumway et al., 2000). Within the field of data mining, such methods are commonly used to quickly obtain approximate solutions for a given problem. To do this, data are compared using major features of the transformations (usually at a much lower dimensionality than the original data) and are subsequently verified either by using original data, or by reversing the transformations (Ye, 2003). This verification step is important since feature selection can entail a loss of information which varies with both the selection process used and the composition of input data. Due to the ordered nature of genetic data, many of these transformation approaches can also be applied to sequences of nucleotides (Silverman and Linsker, 1986) and amino acids (Cheever et al., 1989). Since most of these transformation approaches are suitable only for numerical sequences, an appropriate numerical sequence representation must be applied prior to using these methods.

Many methods for representing symbolic nucleotide sequences as numerical sequences have been proposed (Kwan and Arniker, 2009), permitting the application of various digital signal processing techniques to nucleotide sequences. They include the Voss method (Voss, 1992), the DNA walk (Berger et al., 2004) (Lobry, 1996), the real number (Chakravarthy et al., 2004), complex number (Anastassiou, 2001), and tetrahedron methods (Silverman and Linsker, 1986). Fundamentally, all of these sequence representation approaches convert a nucleotide sequence into a numerical sequence by assigning a number or vector to each nucleotide. Some of the methods, including the complex number and the real number methods, introduce biasing mathematical properties to the transformed sequence, yet are appropriate for use in certain applications such as the detection of AT or GC biases in sequence composition (Arneodo et al., 1998). The Voss and the tetrahedron methods, among others, are notable as they do not introduce biases in internucleotide distance, and thus represent a good starting point for pairwise comparison of nucleotide sequences.

To investigate the applications of time series data mining techniques for sequence alignment, we used three established dimensionality reduction methods to align simulated short DNA sequencing reads both with and without use of a reference sequence. We benchmarked the accuracy of our short read aligner implementation against existing tools, and successfully demonstrated the applicability of our approach to *de novo* assembly.

## 2 METHODS

### 2.1 Symbolic to numeric sequence representations

For the numerical representation step, we primarily focus on the Voss representation (Voss, 1992). This is a fixed mapping approach which turns a nucleotide sequence with *n* dimensions into a four-dimensional binary matrix of length n × 4.

Each row vector in the matrix represents a nucleotide (symbols *G*, *C*, *A* and *T*), while each column vector represents a sequence position. Binary values are assigned to each cell, indicating the presence or absence of a nucleotide at each sequence position (equation 1). *V*_*4i*_ is the binary indicator for presence of a nucleotide in the *i*^*th*^ position of the sequence *S* with *n* nucleotides. Since the Voss representation does not introduce internucleotide mathematical bias, the pairwise distances between all the sets of non-similar transformed nucleotide is the same (for example, the distance between *A* and *T*is equal to the distances between *A* and *C* as well as *A* and *G*).

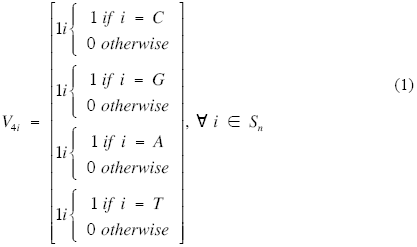

By contrast, a DNA walk assigns trajectories to each sequence nucleotide and may be used to depict genetic sequences in Euclidean space. In two-dimensional DNA walks (Berger et al., 2004), the pyrimidines cytosine and thymine (symbols *C* and *T*) have an upward trajectory while the purines adenine and guanine (*A* and *G*) have a downward trajectory. The trajectory of the sequence extends in a cumulative manner with each consecutive nucleotide. In three-dimensional DNA walks (Lobry, 1996), the cardinal directions north, south, east or west are assigned to each nucleotide.

### 2.2 Sequence Transformation

Effective methods for transforming/approximating sequential data should: *i)* accurately transform/approximate data without loss of useful information, *ii)* have low computational overheads, *iii)* facilitate rapid comparison of data, and *iv)* provide lower bounding—where the distance between data representations is always smaller than or equal to that of the original data—guaranteeing no false negative results (Faloutsos et al., 1994). The DFT and the DWT transformation methods and PAA approximation method satisfy these requirements and are widely used for analyzing discrete signals (Mitsa, 2010); we thus use them here. These methods can be used to transform/approximate nucleotide sequence numerical representations to different levels of resolution, permitting reduced dimensionality sequence analysis (compared in Fig. 1).

**Fig. 1.**
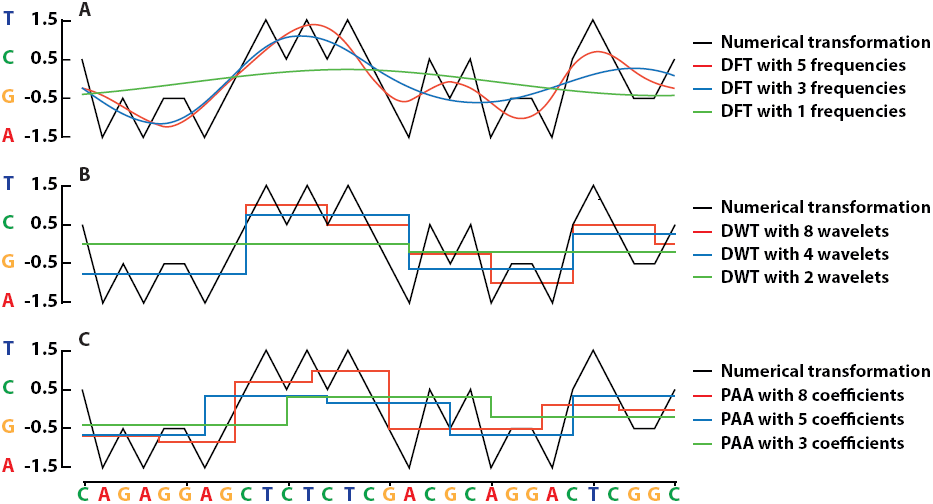

A numerically represented DNA sequence transformed at various levels of spatial resolution using the discrete Fourier transform (DFT) of the whole sequence (A), the Haar discrete wavelet transform (DWT) (B), and piecewise aggregate approximation (PAA) (C). A 30 nucleotide sequence (top) is represented as a numerical sequence (black lines) using the real number representation method (*T*=1.5, *C*=0.5, *G*=-0.5 and *A*=-1.5). DFT approximations of the sequence with 5 (red), 3 (blue) and 1 (green) respective Fourier frequencies (A). DWT approximations of the same sequence with 8 wavelets (red), 4 wavelets (blue), and 2 wavelets (green) (B). PAA approximations of the same sequence with 8 (red), 5 (blue) and 3 (green) respective coefficients (C).

The DFT decomposes a numerically represented nucleotide sequence with *n* positions (dimensions) into a series of *n* frequency components ordered by their frequency. However, since the DFT method has high time complexity, *O(n*^*2*^), the fast Fourier transform (FFT) algorithm (Cooley and Tukey, 1965) with lower time complexity, *O(n log(n))*, is typically used instead. A prerequisite of the FFT algorithm is a signal with length equal to an integer exponent of two, 2^*n*^. Where sequences have a length other than 2^*n*^, they are padded with zeros up to the next integer exponent of two prior to application of the FFT. Because the DFT decomposition of a real signal is conjugate symmetric (Briggs, 1995), half of the resulting frequency components can be safely discarded yet still permit complete signal reconstruction from the remaining frequencies. In time series data mining, a subset of the resulting Fourier frequencies are used to approximate the original sequence in a lower dimensional space (Agrawal et al., 1993), and the tradeoff between analytical speed and accuracy can be varied according to the number of frequencies considered (Mörchen, 2003).

The DWT is a set of averaging and differencing functions that may be used recursively to represent sequential data at different resolutions (Chan and Fu, 1999; Jensen and la Cour-Harbo, 2001). Unlike the DFT, the DWT provides time-frequency localisation, so better accommodates changes in signal frequency over time (non-stationary signals), compared with the DFT and related methods (Wu et al., 2000). As with the FFT, a drawback of the DWT is its requirement of input with length of an integer exponent of two, 2^*n*^. Where sequences have a length other than 2^*n*^, artificial zero padding is therefore added to increase the size of the signal up to the next integer exponent of two prior to application of the DWT. The corresponding DWT transformations are then truncated in order to remove the bias associated with artificial padding (Percival and Walden, 2006). For example, in order to generate the DWT transformation of a time series with 500 data points to a resolution of three, i.e. 2^3^, artificial padding must be added to increase its length to 512 (2^9^) – the next integer exponent of two. In this case, the final, eighth wavelet should be truncated so as to avoid introducing bias.

In PAA a numerical sequence is divided into *n* equally sized windows, the mean values of which together form a compressed sequence representation (Geurts, 2001; Keogh et al., 2001). The selection of *n* determines the resolution of the approximate representation. While PAA is faster and easier to implement than the DFT and the DWT, unlike these two methods it is irreversible, meaning that the original sequence cannot be recovered from the approximation. Fig. 1 depicts examples of the DFT, the DWT and PAA transformations of a short nucleotide sequence.

### 2.3 Similarity search approaches for sequential data

In addition to the DFT, the DWT and PAA, suitable methods for measuring the similarity of sequential data or transformations include the Lp-norms (Yi and Faloutsos, 2000), dynamic time warping (DTW) (Keogh and Ratanamahatana, 2005), longest common subsequence (LCSS) (Vlachos et al., 2002) and alignment approaches such as the Needleman-Wunsch and Smith-Waterman algorithms. Euclidean distance is arguably the most widely used Lp-norm method for sequential data comparison. Lp-norms are straightforward and fast to compute, but require input data of the same dimensionality (sequence length). For comparing sequences of different lengths, a workaround is to simply truncate the longer of the sequences. Furthermore, Lp-norm methods do not accommodate for shifts in the *x*-axis (time or position) and are thus limited in their ability to identify similar features within offset data. Elastic similarity/dissimilarity methods such as LCSS, unbounded DTW and various alignment algorithms permit comparison of data with different dimensions and tolerate shifts in the *x*-axis. These properties of elastic similarity methods can be very useful in the analysis of speech signals, for example, but can be computationally expensive (Kotsakos et al., 2013) in comparison with measures of Euclidean distance. Several approaches have been proposed to permit fast search with DTW, including the introduction of global constraints (wrapping path) or the use of lower bounding techniques such as LB_keogh (Keogh and Ratanamahatana, 2005).

Similarity search strategies can be broadly classified into whole matching and subsequence matching methods. Whole matching is appropriate for comparing sequences with similar overall characteristics—including length—with the queried sequence. Subsequence matching, by contrast, is more suited to identifying similarity between a short query sequence and limited, subsequence regions of longer sequences. Subsequence matching can however be effectively adapted for whole matching purposes through copying to a new dataset each subsequence falling within a sliding window of the longer sequence. The newly created dataset is then used for whole matching indexing similarity search (Das et al., 1998). While using subsequence similarity search for clustering is flawed (Keogh and Lin, 2005), here this approach is only used as a subroutine for extracting all possible subsequences from the genome, which are then used for indexing short reads to the genome. In spite of the storage redundancies associated with this approach, it is both fast and easily implemented, and so was used for our alignment algorithm.

## 3 RESULTS

In order to assess the performance of our sequence transformation and approximation approach, we implemented reference-based and *de novo* read aligners. The four key stages of our approach (Fig. 2) are as follows: *i)* transforming nucleotide sequence characters into numerical sequences, *ii)* creating approximate transformations of reads and, where appropriate, of a reference sequence, *iii)* performing accelerated comparison of the sequence approximations created in the previous step in order to identify candidate alignments, and *iv)* verifying and finishing candidate alignments using original, full-resolution sequences and the Needleman-Wunsch (NW) global alignment algorithm.

**Fig. 2.**
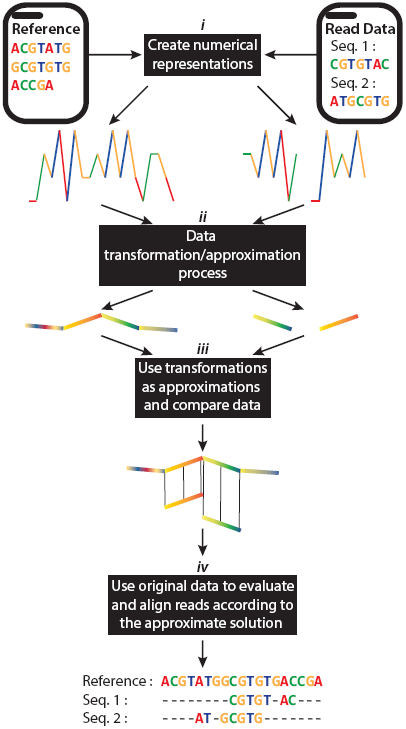

Overview of our short read aligner implemented using time series transformation/approximation methods. *(**i**)* Creation of numerical representations of input sequences. *(**ii**)* Application of an appropriate signal decomposition method to transform sequences into their feature space. *(**iii**)* Use of approximated transformations to perform rapid data analysis in lower dimensional space. *(**iv**)* Validation of inferences against original, full-resolution input sequences. In the case of reference-based alignment, approximated read transformations were compared with a reference sequence. In our *de novo* implementation, pairwise comparisons were performed between all of the approximated read transformations.

### 3.1 Read simulation

To facilitate performance evaluation of our read aligner implementation, sequencing reads were simulated. Using CuReSim (Caboche et al., 2014), sixteen pyrosequencing runs of an HIV-1 HXB2 reference sequence (9719 nucleotides, GenBank accession: K03455.1) were simulated with a mean single read length of 400 nucleotides and mean coverage depth of 100 reads: *i)* in the absence of variation or sequencing error, with *ii)* nucleotide insertion/deletion rates of 1–5%, *iii)* nucleotide substitution rates of 1–5%, and iv) matching insertion/deletion and substitution rates of 1–5% (2%, 4%, 6%, 8%, and 10% overall variation) so as to simulate the heterogeneity present in diverse viral populations. Read quality was simulated at a fixed value of Sanger Q30. CuReSim recorded the exact origin of each simulated read with respect to the reference sequence, enabling later evaluation of alignments using in terms of precision, recall and *F*-Score.

### 3.2 Reference-based alignment

To demonstrate the applicability of sequential data transformations and feature selection in read alignment, we implemented a naive sequential scanning *k*-Nearest Neighbours (*k*NN) read alignment in the Matlab environment. In our implementation (Table 1), the prerequisite numerical sequence representation is performed using a four-dimensional binary mapping approach proposed by Voss, chosen since, as already noted, it introduces no biases in internucleotide distance, and since its binary nature removes the need for normalisation prior to analysis. Transformations of numerically represented sequence *k*-mers are constructed using one of three implemented methods: the DFT, the DWT and PAA. Euclidean distance was used as a similarity measure for read and reference sequence comparison as it is: *i)* fast and easily implemented, *ii)* applicable to many different transformation/approximation methods, and *iii)* its performance compares with more sophisticated ‘elastic’ similarity methods for *k*NN search in medium to large datasets (Wang et al., 2013). After generating candidate alignments between reads and reference *k*-mers through pairwise comparison of their transformations, these candidates alignments are verified NW global alignment of their corresponding original sequences. Finally, NW alignment scores are used to identify best alignments for each read and to reject false positives, and the algorithm’s gapped output is used to construct alignments in the widely used Sequence Alignment/Map (SAM) file format.

**Table 1.**
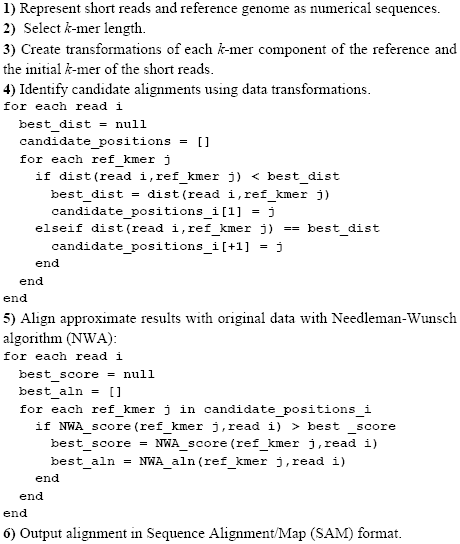
Pseudocode of the alignment procedure.

### 3.3 Benchmarking

CuReSim’s companion tool CuReSimEval was used to quantify alignment accuracy in terms of *F*-score, a balanced measure of precision and recall. The relative performance of three numerical sequence transformation methods was assessed against simulated reads with 6% and 10% variation from the HIV-1 reference sequence. These two simulated datasets were aligned using an otherwise identical implementation using: *i)* the DWT transformation, *ii)* the DFT transformation, *iii)* PAA approximation of the Voss sequence representation of the nucleotide sequences and, finally, *iv)* the full resolution Voss representations of the sequences; thus removing the overhead of building approximate sequence transformations.

We observed that in spite of their associated overheads, the use of approximate sequence transformations dramatically reduced execution time and yielded alignments of equal or greater accuracy than could be obtained using uncompressed sequences. For simulated reads with 6% variation, alignment of uncompressed (baseline) sequences was performed in 734 seconds (s) with an *F*-score of 0.697. For the same dataset, PAA-approximated sequences were aligned fastest and most accurately with an execution time of 54s (~14-fold faster than baseline) and an *F*-score of 0.704, while the for the DWT implementation sequences were aligned in 445s (~2 fold faster) with an *F*-score of 0.698, and the DFT implementation sequences were aligned in 120s (~6-fold faster) with an *F*-score of 0.692.

Alignment of reads with 10% overall variation was performed in 853s using uncompressed sequences giving rise to an *F*-score of 0.567. The PAA approximated sequences were again aligned fastest in 60s (again some 14-fold quicker) with an *F*-score of 0.572, while the DWT transformed sequences were aligned most accurately (*F*-score 0.577) but most slowly with an execution time of 516s, and the DFT transformed sequences were aligned in 138s with an *F*-score of 0.575.

The alignment accuracy of our signal decomposition approach was also evaluated alongside the existing read aligners Bowtie2 (Langmead and Salzberg, 2012), BWA-MEM (Li and Durbin, 2009), Mosaik (Lee et al., 2014), and Segemehl (Otto et al., 2014). Three variants of our approach using the DFT, the DWT and PAA dimensionality reduction methods were tested, using otherwise identical parameters and *k*-mers of length 100–300 nucleotides. Existing tools were all configured with default parameters and run once per dataset, so as to provide a conservative comparison of our implementation performance. Note, the results we present for our approach correspond to the worst performing *k*-mer for each dataset.

Using a strict definition of mapping correctness where reads are considered correctly mapped only if their exact start position is identified, the performance of the tested aligners was relatively similar (Fig. 3A-C). Segemehl generally produced the most accurate alignments in terms of F-score, its accuracy falling behind those of other tools only at the highest rate of sequence variation tested. Our implementation was outperformed by several existing tools in terms of strict start position accuracy. This can mostly be attributed to the use of a global alignment algorithm for the alignment finishing step, which, in the presence of insertion variation near the beginning of reads, tended to insert gaps rather than truncate the aligned region. Consequently, in some cases our approach identified a starting position one or two nucleotides before that deemed correct by CuReSimEval. The challenges associated with assessing alignment correctness for benchmarking purposes are discussed by (Holtgrewe et al., 2011).

**Fig. 3.**
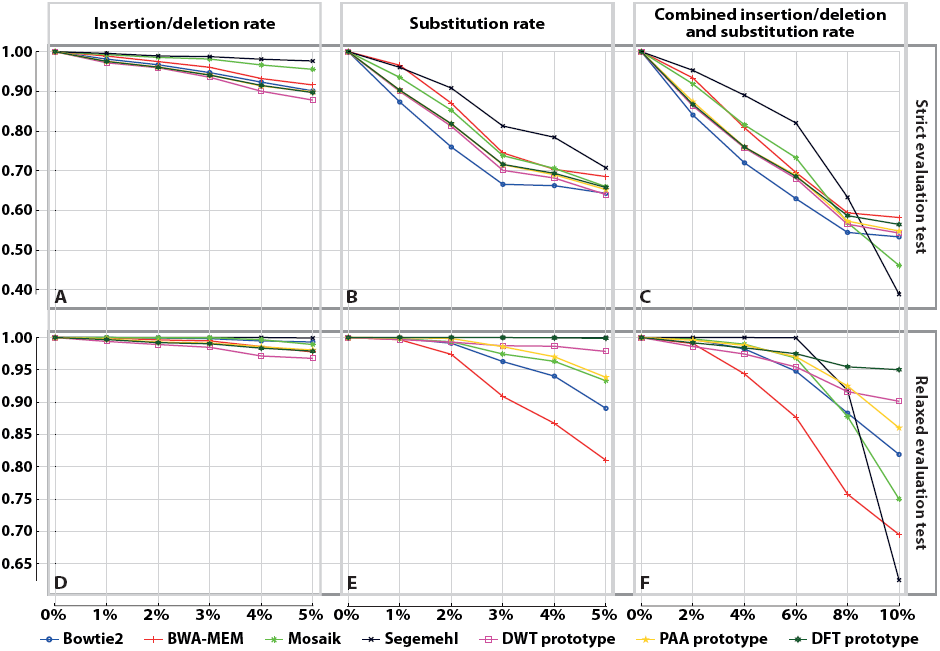
Accuracy of our prototype aligner variants and four established tools in aligning simulated 400 nucleotide (mean length) reads with varying levels of sequence variation. A-C depict alignment accuracies according to a strict criteria, requiring identification of a read's exact starting position determined by the read simulator, while in D-F a relaxed (±10 nucleotides) criterion is used. A and D show results for reads with 0-5% insertion/deletion variation, while B and E correspond to reads with 0-5% substitution variation. C and F show obtained accuracies for reads with combined, equally contributing insertion/deletion and substitution rates of 0-10%.

Accordingly, a relaxed ‘*correctness*’ definition (correct start position ±10 nucleotides) was also considered (Fig. 3D-F), negating the impact of: *i)* multiple possible alignments associated with simulated variation near the start of reads, *ii)* use of different gapped alignment algorithms, and *iii)* the use of different algorithm parameters including scoring matrices and match, mismatch and gap extension penalties. Under this relaxed definition, our signal decomposition approach yielded the most accurate overall alignments for reads containing both insertion/deletion and substitution variation (Fig. 3F), while our DFT-based implementation offered joint best performance for reads containing only substitutions (Fig. 3E). Alignment accuracy for reads containing only insertion/deletion variation was comparable but slightly below the average of existing tools tested (Fig. 3D). Notably, the relative performance of our approach improved considerably with increasing rates of sequence variation, and existing tools were outperformed at the highest rates tested.

### 3.4 *De novo* assembly

To demonstrate the applicability of our approach to the *de novo* assembly of short reads, we implemented a naive algorithm for all-against-all *k*-mer comparison using wavelet transformations (Fig. 4). Reads are first represented as numerical sequences using the Voss method. Every *k*-mer of each numerically represented read is subsequently identified and transformed to lower dimensional space using the DWT method. The *k*-mers’ transformations are then compared with one another to establish their pairwise similarities in terms of Euclidean distance, and to construct a weighted graph (Fig. 4A). Finally, a breadth-first search (BFS) algorithm identifies the shortest path through the graph (Fig. 4B), and after attribution of *k*-mers to their corresponding reads, yields an assembly of short reads (Fig. 4C). Additionally, a numerical representation such as the DNA walk may subsequently be used to aid visualisation of the assembly (Fig. 4D). 256mers were used in our tests, and each 256-mer representation was approximated to a length of 16.

**Fig. 4.**
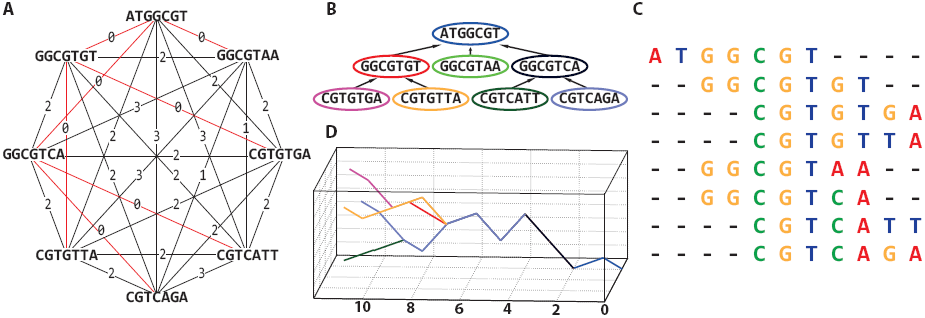

A *de novo* read assembly methodology for numerically represented nucleotide sequences. All-against-all sequence comparison (A) enables construction of a read graph with weighted edges. The weight assigned to each edge is the smallest pairwise distance between every possible *k*-mer representation of the two reads. The shortest path in the graph is identified with a breadth-first search algorithm (B), thereby enabling read alignment (C). A DNA walk of aligned reads (D) may subsequently be used as a three-dimensional graphical portrayal of the reads, illustrating alignment characteristics.

We applied our *de novo* aligner algorithm to the assembly of simulated short read data from viral populations (HIV-1). The dimensionality of the numerical sequence representations was reduced by ~16-fold using the DWT prior to alignment, and the deviation of the resulting assemblies from the reference sequence used for read simulation was quantified using CuReSimEval. Fig. 5A illustrates the three-dimensional ‘walk’ of the HIV-1 reference sequence HXB2, while Fig. 5B, 5C and 5D depict the three-dimensional surface plots, of *de novo* alignments for reads with 2%, 4% and 6% variation. The alignments of the 0%, 2%, 4% and 6% variation datasets had *F*-scores of 1, 0.9979, 0.9255 and 0.7772 respectively. Despite the high levels of variation in the data, and the use of large *k*-mers, our approach yields accurate alignments, highlighting the potential benefits of our approach for *de novo* alignment for processing nucleotide sequences.

**Fig. 5.**
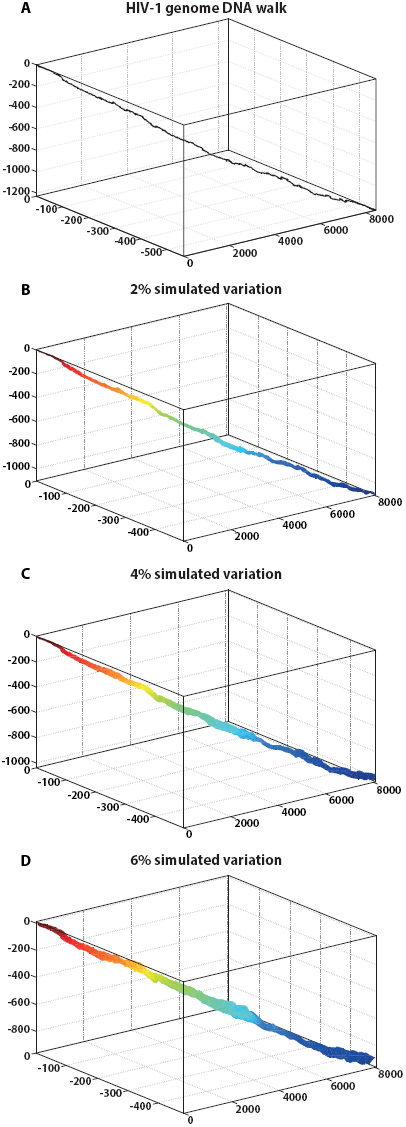

A three-dimensional DNA walk of the HIV-1 HXB2 genome (A) also plotted with *de novo* alignments of three simulated HXB2 sequencing datasets of 2%, 4%, and 6% rates of combined insertion/deletion and substitution variation (B-D respectively). Plotting DNA walks of aligned short reads enables intuitive visualisation of the nature and extent of sequence diversity across a genomic region, with sequence variants each represented by a distinct trajectory through space.

## 4 DISCUSSION

Data compression methods enabling reversible compression of one-dimensional and multivariate signals, images, text and binary data are well established (Hendriks et al., 2013; Sheybani, 2011; Tapinos and Mendes, 2013). Surprisingly, these methods have seen limited application to the problem of nucleotide sequence alignment. Here we have demonstrated the application of a flexible sequence alignment heuristic leveraging established signal compression and decomposition methods. Our implementation aligned simulated viral reads with comparable overall precision and recall to existing tools, and excelled in the alignment of reads with high levels of sequence diversity, as often observed in RNA virus populations (Archer et al., 2010). Our results show that full, nucleotide-level sequence resolution is not a prerequisite of accurate sequence alignment, and that analytical performance can be preserved or even enhanced through appropriate dimensionality reduction (compression) of sequences. For example, six-fold and fourteen-fold reductions in execution time were observed during our tests of the DFT and PAA-represented sequences, yet in both cases alignment accuracy was marginally better than that obtained using full resolution sequences. The approach’s applicability to *de novo* assembly of divergent sequences was also demonstrated. While our implementation makes use of *k*-mers, the nature of the transformation/compression approaches used means that optimal *k*-mer selection is considerably less important than it is with conventional exact *k*-mer matching methods. The inherent error tolerance of the approach also permits the use of larger *k* values than normally used with conventional sequence comparison algorithms, reducing the computational burden of pairwise comparison, and thus, in *de novo* assembly specifically, the complexity of building and searching an assembly graph.

Efficient mining of terabase scale biological sequence datasets requires us to look beyond substring-indexing algorithms towards more versatile methods of compression for both data storage and analysis. The use of probabilistic data structures can reduce considerably the computer memory required for in-memory sequence lookups at the expense of a few false positives, and Bloom filters and related data structures have seen broad application in *k*-mer centric tasks such as error correction (Shi et al., 2010), *in silico* read normalisation (Zhang et al., 2014) and *de novo* assembly (Berlin et al., 2014; Salikhov et al., 2013). However, while these hash-based approaches perform very well on datasets with high sequence redundancy, for large datasets with many distinct *k*-mers, large amounts of memory are still necessary (Zhang et al., 2014). Lower bounding transformations and approximation methods (such as the DFT, the DWT and PAA) exhibit the same attractive one-sided error offered by these probabilistic data structures, yet—unlike hashing—construct intrinsically useful sequence representations, permitting their comparison with one another. Furthermore, transformations allow compression of standalone sequence composition, enabling flexible reduction of sequence resolution according to analytical requirements, so that redundant sequence precision need not hinder analysis. In large datasets, the associated reductions in resource usage can be significant. While the problem of read alignment to a known reference sequence is largely solved, the assembly of large and/or poorly characterised sequenced genomes remains limited by computational methods. Moreover, consideration of the metagenomic composition of mixed biological samples further extends the scope and scale of the assembly problem beyond what is tractable using conventional sequence comparison approaches. Through implementing of reference-based and *de novo* aligners, we have demonstrated that the analysis of compressed representations provide a tractable, memory-efficient and versatile approach to the short-read-based reconstruction of genomes and metagenomes.

In conclusion, short nucleotide sequences may be effectively represented as numerical series, enabling the application of existing analytical methods from a variety of mathematical and engineering fields for the purposes of sequence alignment and assembly. By using established signal decomposition methods, it is possible to create compressed representations of nucleotide sequences, permitting substantial reductions in the spatiotemporal complexity of their analysis without necessarily compromising analytical accuracy.

## ACKNOWLEDGMENTS

We thank Mattia Prosperi for helpful discussion. Funding: AT has been supported by the Wellcome Trust [097820/Z/11/B] and BBSRC [BB/H012419/1 & BB/M001121/1], and BC by a BBSRC DTP studentship.

